# Dynamic network regulating phosphatidic acid homeostasis revealed using membrane editing coupled to proximity labeling

**DOI:** 10.1101/2024.09.14.612979

**Authors:** Reika Tei, Jeremy M. Baskin

## Abstract

Cellular lipid metabolism is subject to strong homeostatic regulation, but players involved in and mechanisms underlying these pathways remain mostly uncharacterized. Here, we develop and exploit a “Feeding–Fishing” approach coupling membrane editing using optogenetic lipid-modifying enzymes (feeding) with organelle membrane proteomics via proximity labeling (fishing) to elucidate molecular players and pathways involved in homeostasis of phosphatidic acid (PA), a multifunctional lipid central to glycerolipid metabolism. By performing proximity biotinylation using a membrane-tethered TurboID alongside membrane editing to selectively deliver phosphatidic acid to the same membrane, we identified numerous PA-metabolizing enzymes and lipid transfer proteins enriched in and depleted from PA-fed membranes. Subsequent mechanistic analysis established that PA homeostasis in the cytosolic leaflets of the plasma membrane and of lysosomes is governed by a select subset of PA metabolic pathways and, via divergent molecular mechanisms, several members of the lipid transfer protein superfamily capable of mediating interorganelle lipid transport. More broadly, the interfacing of membrane editing with organelle membrane proteomics using proximity labeling represents a powerful and generalizable strategy for revealing mechanisms governing lipid homeostasis.

## Introduction

Membranes are complex and dynamic assemblies that regulate myriad cellular processes. The lipid composition of membranes, crucial to membrane biophysical properties and the functions of the membrane-associated proteome, is determined by a balance of activities of lipid-metabolizing enzymes and lipid transfer proteins (LTPs). Efforts to decode these lipid–protein networks are complicated by the rapid diffusion, trafficking, and, in many cases, short metabolic half-lives of lipids.

One type of lipid emblematic of these challenges is phosphatidic acid (PA)^1^. A central player in lipid metabolism, PA is produced by at least four different biosynthetic routes and metabolized into phospholipids, triglycerides, or fatty acids by three pathways^2,3^. Though only 1% of the lipidome, PA is present in most organelle membranes, and its levels transiently rise downstream of signaling from numerous cell-surface receptors and soluble signaling enzymes^4,5^. Because such PA bursts induce potent signaling events, PA is subject to strong homeostatic regulation, with its levels tightly controlled in space and time by modulation of both biosynthetic flux and regulated interorganelle transport by LTPs^4–7^. Though several PA-metabolizing enzymes and PA-specific LTPs have been characterized, it is not well understood how cells sense changes to levels of PA to regulate the activities of these enzymes and transporters to ultimately restore PA levels and achieve homeostasis.

Membrane editing is an emerging strategy to precisely manipulate the lipid composition of organelle membranes within living cells to study physiological functions of individual lipids. We have previously devoted substantial efforts toward development of membrane editors for PA. First, we designed an optogenetic phospholipase D (optoPLD) to acutely produce PA on target organelle membranes^8^. PLDs produce PA by hydrolysis of abundant phospholipids, and optoPLD uses a light-inducible heterodimerization system to recruit PLD onto desired membranes. Moreover, the ability of PLDs to additionally catalyze transphosphatidylation with exogenously supplied primary alcohols to produce a variety of natural and unnatural phospholipids underscores the versatility of optoPLD for editing the phospholipidome^9^.

To overcome the relatively modest activity of the first-generation optoPLD, we greatly improved its performance by using directed evolution in mammalian cells. The resultant super-active optoPLDs (superPLDs), which acquired several mutations that optimized performance of this disulfide-containing, secreted protein from *Streptomyces sp.* PMF in the mammalian intracellular environment, exhibited up to 100-fold higher activity in cells compared to the original editor (wild-type optoPLD). SuperPLDs were demonstrated as potent membrane editors for spatiotemporally defined editing of phospholipids with organelle-level precision in live cells.

Here, we exploit a “Feeding–Fishing” strategy to map such lipid–protein networks and reveal mechanisms of how cells respond to changes in the lipid composition of their organelle membranes. In this approach, we combined superPLD-mediated membrane editing with proximity-dependent biotinylation using a membrane-tethered TurboID, followed by tandem mass tagging (TMT)-based mass spectrometry (MS) to enable protein identification and quantification. In this manner, “feeding” PA to a membrane and “fishing” out proteins associated with that membrane would identify proteins that accumulate on or are depleted from a target membrane in response to changes in PA levels within that membrane.

Our studies reveal changes to the membrane recruitment of several PA-metabolizing enzymes and LTPs in PA-fed membranes that collectively facilitate removal of excess PA. Analysis of lipid-metabolizing enzymes implicates a select subset of PA metabolic pathways for clearance of PA pools from the plasma membrane and lysosome. Mechanistic investigations using confocal microscopy to visualize PA pools and lipidomics to track changes to the phospholipidome revealed divergent roles for several LTPs recruited to PA-fed membranes in affecting PA metabolism. In particular, we found that Nir2, an LTP known to transport PA between the plasma membrane and ER, could clear PA pools not only from the plasma membrane but also from lysosomes, and SCP2, a broad-spectrum LTP implicated in sterol and fatty acid transport, can also mediate PA clearance. We additionally point to unexpected roles for members of the SMP and ORD domain-containing LTP families in PA homeostasis, including PDZD8, TEX2, and ORP1L. Collectively, membrane editing with superPLD coupled to proximity labeling enables identification of regulators of PA homeostasis, revealing cellular mechanisms underlying rapid PA metabolism and interorganelle transport.

## Results

### Design of a Feeding–Fishing strategy for in situ identification of PA-associated proteins

Our Feeding–Fishing strategy to identify proteins associated with PA-fed membranes involves superPLD-enabled membrane editing coupled to proximity proteomics (**Fig. 1a**). Currently, there are two widely used types of proximity labeling enzymes with distinct labeling mechanisms: promiscuous biotin ligase (e.g., TurboID^10^) and engineered peroxidase (e.g., APEX2^11^). Promiscuous biotin ligase releases a biotin-AMP intermediate that reacts with lysine residues on proximal proteins, whereas proximity labeling with engineered ascorbate peroxidase is mediated by biotin-phenoxyl radicals that react with tyrosine residues^12^. These enzymes are generally expressed in intracellular compartments, where the reactive intermediates are highly confined, and their cross-membrane diffusion is rare.

**Figure 1.**
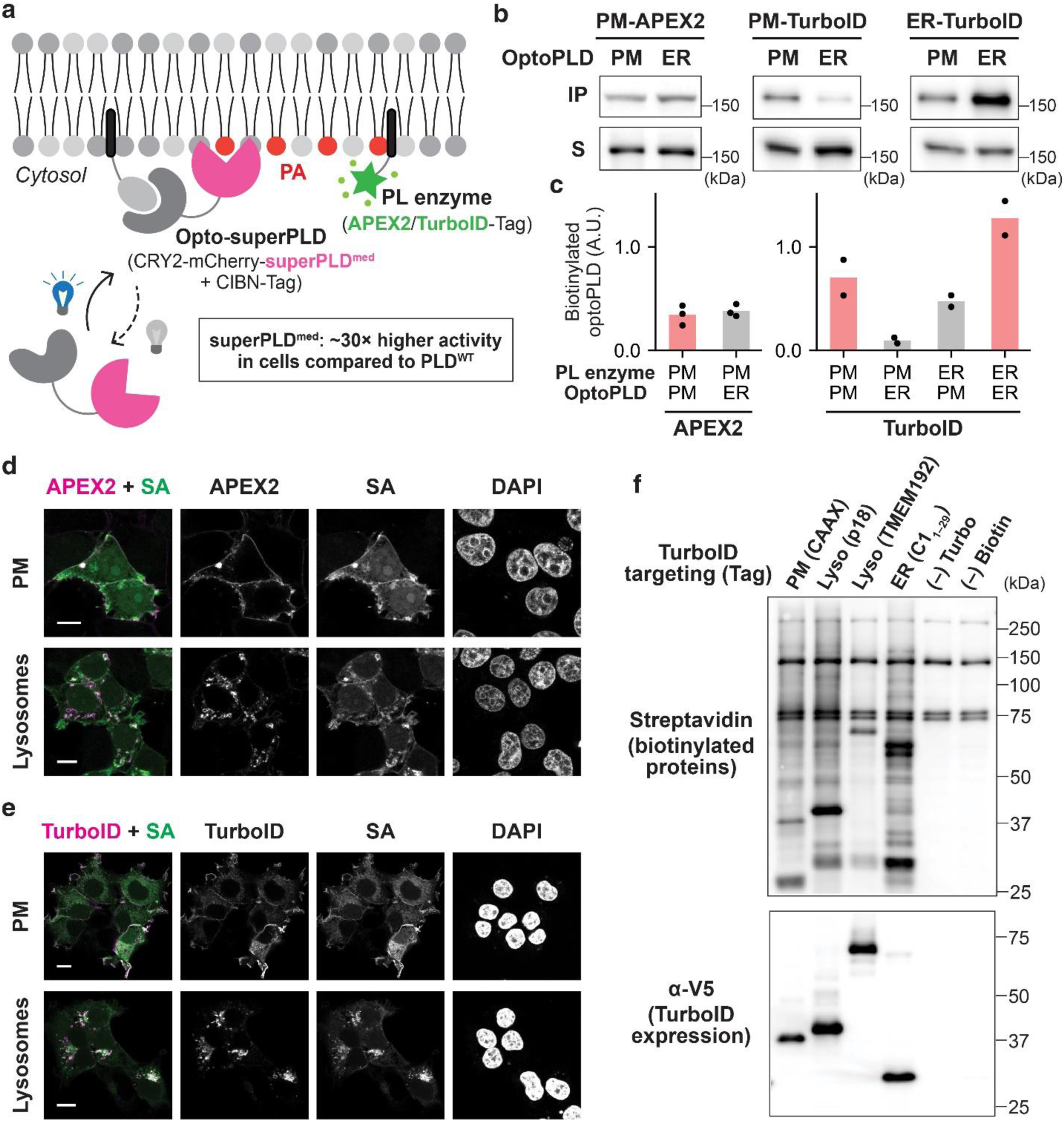
Design of the Feeding–Fishing strategy and selection of optimal proximity labeling enzyme. **a,** Schematic depicting the design of the Feeding–Fishing strategy that couples membrane editing and proximity labeling (PL) for elucidation of regulators of PA homeostasis on different organelle membranes. An optogenetic superPLD (optoPLD) is targeted to a membrane of interest to “feed” it with phosphatidic acid (PA), and a PL enzyme is anchored to the same membrane to enable biotinylation and subsequent “fishing” out of proteins differentially recruited to the lipid-edited membrane. **b,** Comparison of APEX2 and TurboID for proximal organelle membrane proteomics. Shown are representative Western blots evaluating the ability of APEX2 and TurboID to selectively label proteins associated with the membrane of interest. The PL enzyme and optoPLD were targeted to the same vs. different membranes. APEX2 labeling was performed by pre-incubating cells for 30 min with 500 μM biotin-phenol, followed by 1 min with 1 mM H_2_O_2_. TurboID labeling was performed by incubating cells with 500 μM biotin for 10 min.

Because there are limited examples of tethering these enzymes to membranes to map membrane-proximal proteomes, we first explored the feasibility of using these two types of proximity labeling systems in this application, i.e., to selectively label proteins that reside on the same membrane they are tagged to. We expressed APEX2 or TurboID fused to a plasma membrane targeting sequence, Lyn_10_, to localize the enzyme to the cytoplasmic leaflet of the plasma membrane. In the same cells, we co-expressed the wild-type optoPLD (from a bicistronic expression vector, CRY2-mCherry-PLD-P2A-CIBN-Tag where Tag is an organelle-targeting sequence) targeted to the cytosolic leaflet of either the plasma membrane or endoplasmic reticulum (ER) membrane. Surprisingly, plasma membrane-tagged APEX2 showed no difference in the labeling on optoPLD on the same (plasma membrane) vs. different (ER) membrane, whereas TurboID preferentially labeled optoPLD on the same membrane (**Fig. 1b–c**). Consistent with this result, TurboID, when targeted to specific membranes, exhibited better colocalization with biotinylated proteins than APEX2 (**Fig. 1d–e**). From these results, we concluded that TurboID would be an optimal enzyme for mapping proteins associated with membranes of interest.

Among various membranes that could be targeted by superPLD and TurboID, we selected the cytoplasmic leaflets of the plasma membrane and lysosomal membrane, due to the proposed localization and physiological functions of mammalian PLD1/2, which can initiate PA-based signaling on these membranes^13–15^. We have previously shown that CIBN fused to the CAAX domain of K-Ras^16^ (CIBN-CAAX) and the lysosome-targeting sequence of p18/LAMTOR1^17^ (p18-CIBN) can mediate effective superPLD recruitment and consequent PA enrichment on plasma membrane and lysosomes, respectively^18^. Using the same sequences for targeting TurboID to these membranes, we confirmed that these membrane-tethered TurboIDs showed equally high and promiscuous labeling activity as the originally reported ER membrane-tethered TurboID^10^ (**Fig. 1f**).

Biotinylated proteins were isolated by streptavidin–agarose pulldown and analyzed by Western blot. PM, plasma membrane; ER, endoplasmic reticulum. **c,** Quantification of biotinylated optoPLD in b. n=3 for APEX2 and n=2 for TurboID, where n indicates independent biological replicates. **d–e,** Confocal images of HEK 293T cells expressing APEX2 (d) or TurboID (e) targeted to plasma membrane (PM) or lysosomes. APEX2 and TurboID labeling was performed as described in b, and biotinylated proteins were immunostained using streptavidin–488. Scale bar; 10 μm. **f,** Western blots of cells expressing TurboID targeted to either the PM (using the CAAX domain of K-Ras^16^), lysosomes (using lysosomal-targeting sequence of p18/LAMTOR1^17^ or the full sequence of TMEM192^19^), or the ER (using first 29 amino acids of cytochrome P450 2C1^20^). Cells were treated with 500 μM biotin for 10 min, and cell lysates were analyzed by Western blot to detect biotinylated proteins and TurboID expression.

### Efficient co-expression of Feeding–Fishing components by lentiviral transduction

The Feeding–Fishing strategy requires co-expression of both TurboID and superPLD in cells at high efficiency. To avoid potential long-term effects of chronic expression, we performed transient expression by one-shot, lentivirus-based transduction. Although optoPLD/superPLD is typically expressed using a P2A self-cleavable peptide to ensure equimolar expression of CRY2- mCherry-PLD and CIBN-Tag, we found that for this application, separating these two components enabled more efficient transduction because of the limited size of DNA inserts that each lentivirus can encapsulate^21^. Therefore, our optimized protocol involved a triple lentivirus transduction system with one lentivirus each harboring TurboID-Tag, CRY2-mCherry-PLD, or CIBN-Tag (**Fig. 2a**). To facilitate the incorporation of lentivirus into cells, we used spinfection, which is a method to apply mild centrifugation force during transduction^22^. The ratio of three lentivirus strains was optimized, based on their titers, to achieve the highest co-expression efficiency while keeping cell viability unaffected. With these modifications, we achieved high efficiency (>75%) for transient co-transduction of HEK 293T cells with TurboID and superPLD targeted to either the plasma membrane or lysosomes (**Fig. 2b–c**).

**Figure 2.**
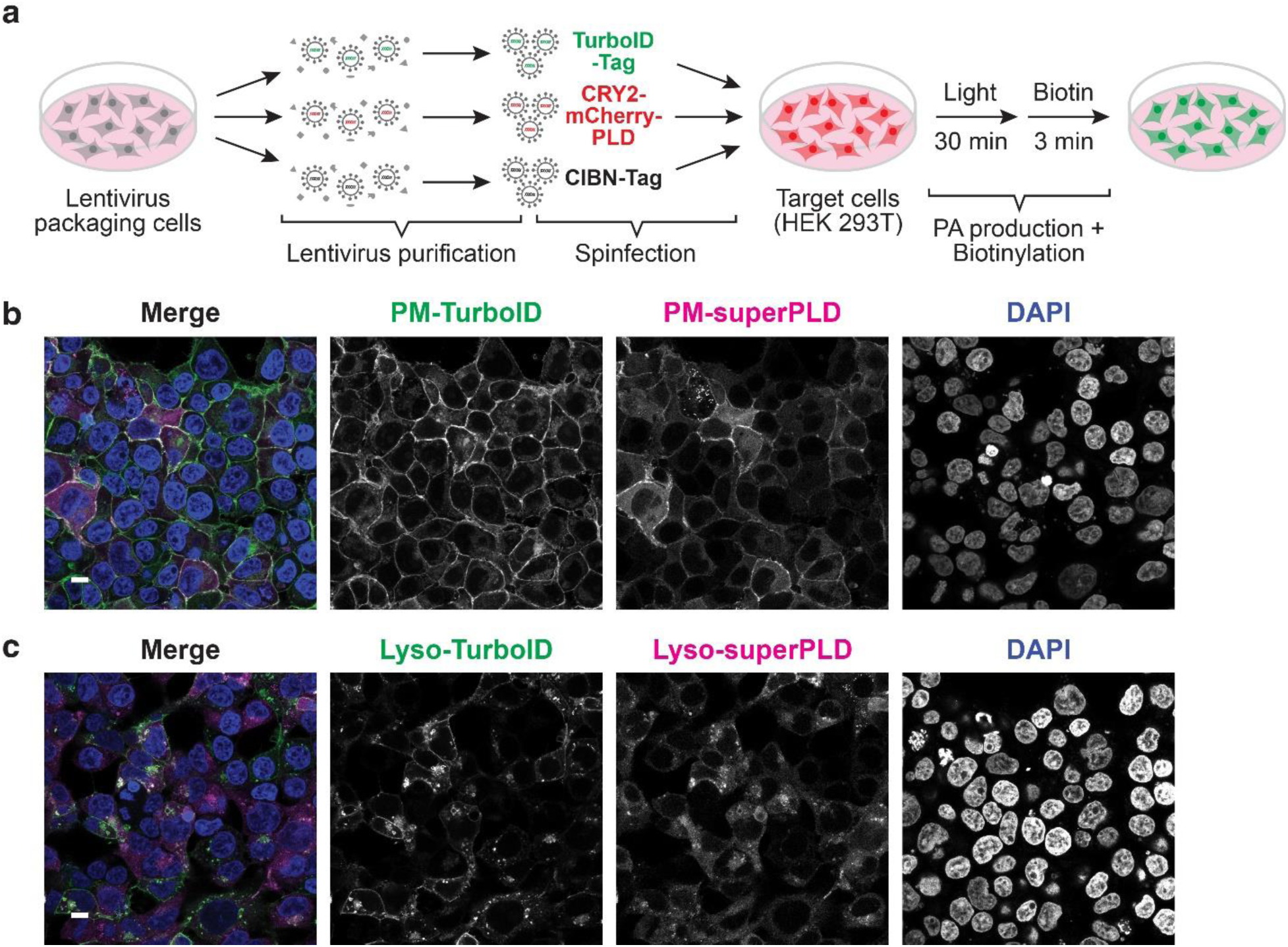
Co-expression of membrane-targeted TurboID and superPLD via one-shot transduction establishes system for Feeding–Fishing studies. **a,** Schematic depiction of transduction strategy to enable efficient and transient co-expression of the three components necessary for Feeding–Fishing proteomics. Lentivirus particles were produced in packaging cells (HEK 293TN) and concentrated by ultracentrifugation. The concentrated virus samples encoding TurboID-Tag, CRY2-mCherry-PLD, or CIBN-Tag, were mixed and used for transduction of HEK 293T cells by spinfection. 48 h after spinfection, the cells were illuminated with blue light for 30 min to stimulate PA production, followed by 150 μM biotin for 3 min. **b–c,** Confocal images of HEK 293T cells co-expressing TurboID and superPLD targeted to plasma membrane (b; PM) or lysosomes (c; Lyso) labeled as in (a), followed by fixation and immunostaining for TurboID (V5) and biotin (streptavidin-Alexa Fluor 488 conjugate). Scale bar; 10 μm.

### Several lipid-metabolizing enzymes and transporters are enriched on PA-fed membranes

We next performed proximity proteomics to identify the effects of local PA production on the proteomes of different membranes. We treated HEK 293T cells co-expressing TurboID and superPLD, targeted to either the plasma membrane or lysosomes, for 30 min with intermittent blue light (5 s every 1 min) to activate superPLD, followed by a 3 min labeling with 150 μM biotin (**Fig. 2a**). As a negative control, we used cells transduced with TurboID and a catalytically dead superPLD (H167A/H440A, deadPLD) that does not produce PA. Biotinylated proteins from two conditions (superPLD vs. deadPLD) and three replicates were combined for TMT 16-plex proteomics analysis to identify proteins that were selectively enriched on or depleted from PA-fed membranes (**Fig. 3a**).

**Figure 3.**
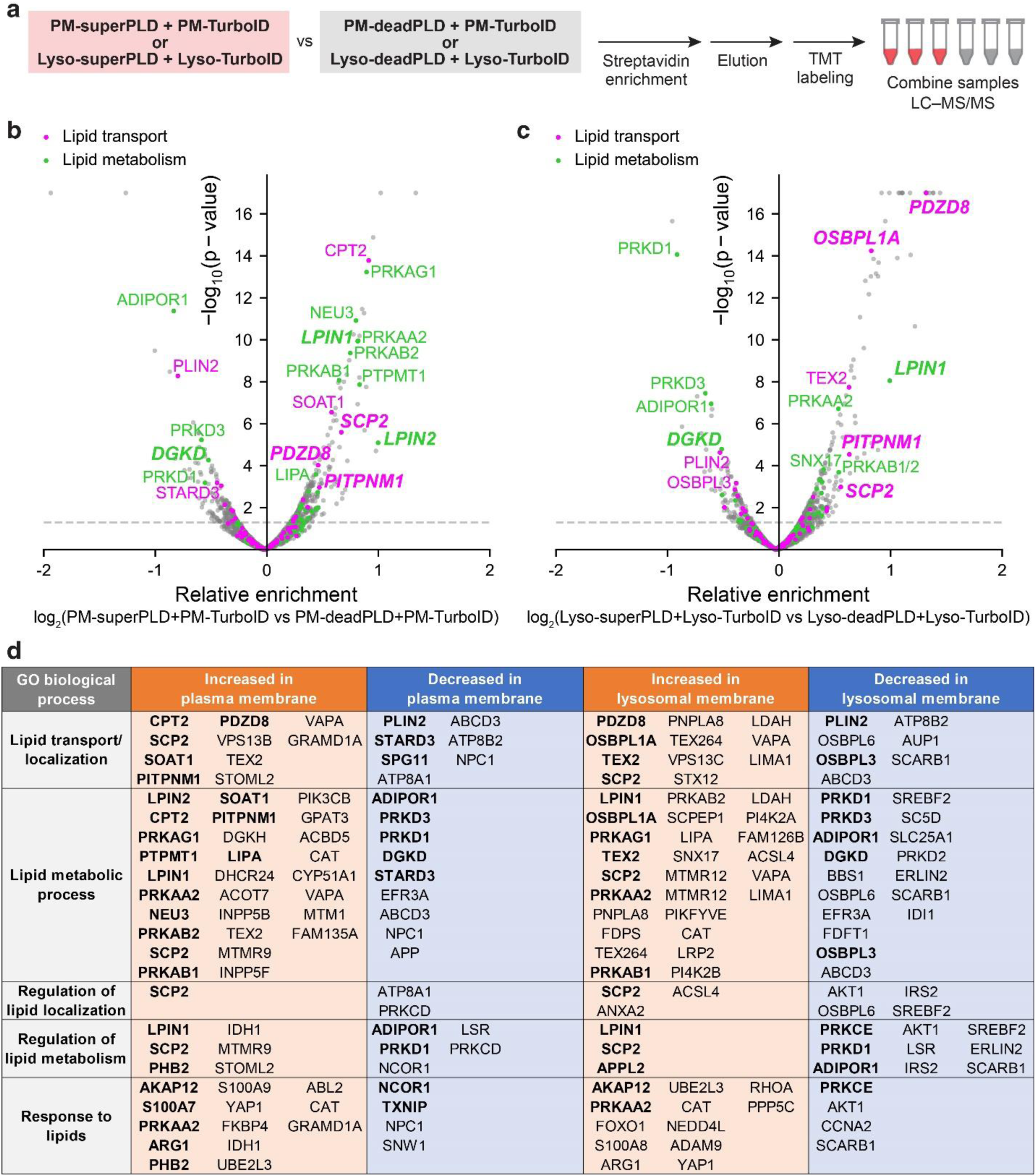
Feeding–Fishing proximity proteomics identifies numerous lipid-related proteins enriched on and depleted from PA-fed membranes. **a,** Schematic of Feeding–Fishing proximity proteomics workflow. Biotinylated proteins from HEK 293T cells co-expressing TurboID and either superPLD or deadPLD targeted to the same membrane (either plasma membrane or lysosomes) were enriched by streptavidin–agarose pulldown, eluted from resin, and subjected to TMT labeling and multiplexed proteomics. **b–c,** Volcano plots showing differential enrichment of proteins on PA-fed vs. PA-unfed plasma membranes (b) or lysosomes (c). Proteins with known functions in lipid transport and lipid metabolism are colored in magenta and green, respectively, and proteins that were further functionally characterized in this study are indicated in bold italic. **d,** Table summarizing proteins that showed significance (abundance ratio *p*-value < 0.05) in enrichment or depletion on PA fed vs. unfed plasma membranes or lysosomes. Protein hits are ordered by fold enrichment, and the hits with particularly high significance (abundance ratio *p*-value < 0.001) are shown in bold. Proteins showing increases (orange in table) are those on the right side of the corresponding volcano plot, whereas those showing decreases (blue in table) are those on the left side of the corresponding volcano plot.

In these Feeding–Fishing proteomics experiments, we detected 4,401 and 4,260 biotinylated proteins from the plasma membrane and lysosomes, respectively, with high correlations between replicates (**Supplementary Tables 1–2**). To identify proteins that were enriched on or depleted from PA-fed membranes, we calculated protein enrichment in the superPLD samples relative to deadPLD controls and plotted abundance ratios and associated *p*-values (**Fig. 3b–c**). Among the hits that showed a significant enrichment on PA-fed membranes (based on the abundance ratio *p*-value) were two known PA-metabolizing enzymes, Lipin-1 (LPIN1) and Lipin-2 (LPIN2), which degrade PA into diacylglycerol (DAG). Correspondingly, DAG kinase delta (DGKD), which produces PA from DAG, was found to be depleted from PA-fed membranes.

Further, the best characterized mammalian PA-transfer protein, PITPNM1 (Nir2), was also enriched on both plasma membranes and lysosomes upon PA production, supporting the validity of the Feeding–Fishing proteomics. We then used the GO biological process annotation to comprehensively highlight proteins associated with lipid biology: lipid localization (which includes lipid transport), lipid metabolic process, regulation of lipid localization, regulation of lipid metabolism, and response to lipids. For the plasma membrane Feeding–Fishing studies, 42 proteins associated with these terms were enriched on this membrane and 18 were depleted; similarly, for lysosomal Feeding–Fishing experiments, 40 such proteins were enriched and 25 were depleted from lysosomal membranes (**Fig. 3d**; also **Supplementary Table 3**).

### Nir2 reduces PA enrichment on plasma membrane as well as lysosomes

PITPNM1/Nir2 is a well characterized PA transfer protein that translocates to ER–plasma membrane contact sites upon PA production (e.g., following phospholipase C activation) and facilitates PA transfer from the plasma membrane to the ER^6^. Because our Feeding–Fishing proteomics detected Nir2 to be enriched not only at the plasma membrane but also on lysosomes upon PA feeding, we overexpressed an miRFP-tagged Nir2 construct to monitor its localization in mammalian cells. miRFP-Nir2 exhibited mostly cytosolic but weakly ER-associated localization under basal conditions, in agreement with a previous study using GFP-Nir2^6^. Upon light-induced superPLD recruitment to the plasma membrane or to lysosomes, miRFP-Nir2 relocalized to these membranes, strongly colocalizing with superPLD (**Fig. 4a–b**). In control cells where deadPLD was used, miRFP-Nir2 did not show any change in its localization.

**Figure 4.**
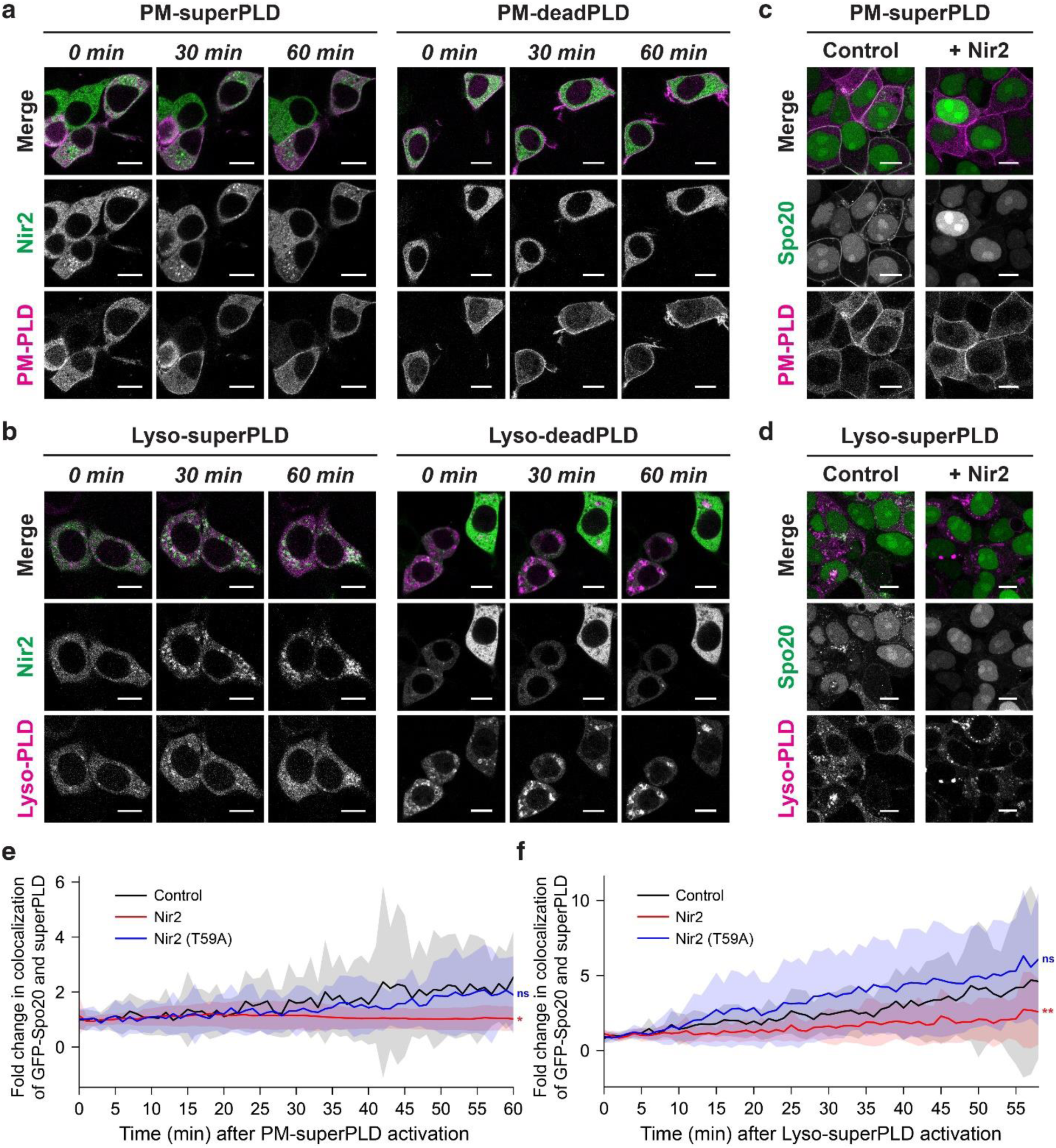
Nir2 is recruited to PA-enriched membranes and reduces local PA enrichment. **a–b,** Confocal microscopy images of HEK 293T cells co-expressing miRFP-Nir2 and either superPLD or deadPLD targeted to the plasma membrane (a, PM) or lysosomes (b, Lyso). Shown are images acquired 0, 30, and 60 min following superPLD or deadPLD recruitment induced by intermittent blue light illumination (5 s per 1 min). **c–d**, Confocal microscopy images of HEK 293T cells co-expressing a PA-binding probe GFP-Spo20 and superPLD targeted to plasma membrane (c; PM) or lysosomes (d; Lyso), without (Control; left panels) or with (+ Nir2; right panels) stable expression of V5-Nir2. Shown are images acquired 60 min following superPLD activation by intermittent blue light illumination (5 s per 1 min). Scale bars, 10 μm. **e–f,** Quantification of imaging data from (c) and (d), measuring colocalization between GFP-Spo20 and superPLD targeted to the plasma membrane (e, PM) or lysosomes (f, Lyso) over the course of the 60 min of superPLD activation. Each graph shows the colocalization timecourse in the cells with empty vector (Control; black), V5-Nir2 (Nir2; red), or a mutant form of V5-Nir2 (T59A) deficient in lipid transfer (Nir2*; blue). Statistical analysis was performed using repeated measures ANOVA (e: *n* = 34 cells, *p* = 0.0032 for Nir2 and 0.1949 for T59A; f: *n* = 20 cells, *p* = 0.0315 for Nir2 and 0.2021 for T59A). Three independent experiments were performed with similar results.

We then tested if Nir2 could exert its PA transfer function on these PA-fed membranes. To minimize any potential interference on its function caused by the fluorescent protein fusion, we replaced miRFP with a V5 tag (GKPIPNPLLGLDST) and used lentiviral transduction to generate HEK 293T cells stably expressing V5-Nir2 (**Fig. S1**). Immunostaining for the V5 tag revealed mostly cytosolic/weak ER localization of V5-Nir2, similar to miRFP-Nir2. To monitor PA localization in these cells, we co-expressed superPLD with the PA reporter GFP-Spo20^23^. GFP-Spo20 was chosen over GFP-PASS, a version designed with a nuclear export signal for improved sensitivity^24^, because the predominantly nuclear localization and lack of cytosolic signal of GFP-Spo20 simplified the quantitative colocalization analysis.

In control wild-type cells, superPLD targeted either to the plasma membrane or to lysosomes triggered PA enrichment on these membranes, as illustrated by accumulation of GFP-Spo20 at the membranes where superPLD was localized (**Fig. 4c**). By contrast, in cells stably expressing V5-Nir2, this PA enrichment did not occur. Quantification of the overlap between GFP-Spo20 and superPLD fluorescence upon superPLD activation revealed little colocalization in V5-Nir2-expressing cells but increasing colocalization over time in cells expressing either the T59A lipid-transfer mutant of Nir2 or empty vector, as continuous superPLD activation would be expected to elevate PA levels (**Fig. 4e–f**). These studies validate not only the ability of the Feeding–Fishing approach to identify a bona fide PA transport protein but also that examining GFP-Spo20 localization upon activation of superPLD in cells expressing a PA transport protein is a straightforward strategy to characterize PA transport proteins.

### The lipid transporter SCP2 reduces local enrichment of PA

We then set out to discover new PA transport proteins from among the Feeding–Fishing proteomics hits (**Fig. 3d**). We selected three additional LTPs — SCP2, PDZD8, and OSBPL1A/ORP1L — that were significantly enriched on PA-fed membranes. Among them, SCP2 and PDZD8 were enriched on both PA-fed plasma membrane and lysosomes, whereas ORP1L was enriched only on PA-fed lysosomes. As we had done with Nir2, we produced HEK 293T cells stably expressing V5-tagged versions of each LTP (**Fig. S1**). Consistent with previous reports, PDZD8-V5 localized to the ER and lysosomes^25^ and V5-ORP1L localized to lysosomes^26^.

V5-SCP2 localized exclusively to punctate structures that did not colocalize with plasma membrane nor lysosomes. Expression of the SCP2 gene leads to two protein isoforms, a full-length 58 kDa SCP2 (also known as SCP-x) and a precursor 15 kDa polypeptide, proSCP2, and both of these proteins undergo post-translational cleavage to produce a 13 kDa C-terminal fragment known as mature SCP2 (mSCP2)^27^. The N-terminal remnant of full-length SCP2/SCP-x, after proteolytic cleavage in the peroxisomes, becomes a functional peroxisomal thiolase^28^, which explains the punctate localization of our V5-SCP2 construct. In line with previous studies^29^, a V5 fusion to mSCP2 showed increased cytosolic localization with much weaker association with peroxisomes (**Fig. S1**).

We then performed the PA enrichment analysis in HEK 293T cells co-expressing GFP-Spo20 and a superPLD recruited to either plasma membrane or lysosomes, as above. Interestingly, HEK 293T cells stably expressing SCP2/SCP-x or proSCP2 showed a significant delay in the colocalization between GFP-Spo20 and superPLD either membrane, indicating reduced PA enrichment on these membranes (**Fig. 5a–d**). The PA reduction effect with proSCP2 was stronger than with SCP2/SCP-x, which aligns with differences in their maturation efficiencies, as cleavage of SCP-x is partial whereas proSCP2 undergoes complete cleavage^27,30^. These studies suggest that SCP2, characterized as a transporter of fatty acids and several other types of phospholipids, may also function as a PA transfer protein.

**Figure 5.**
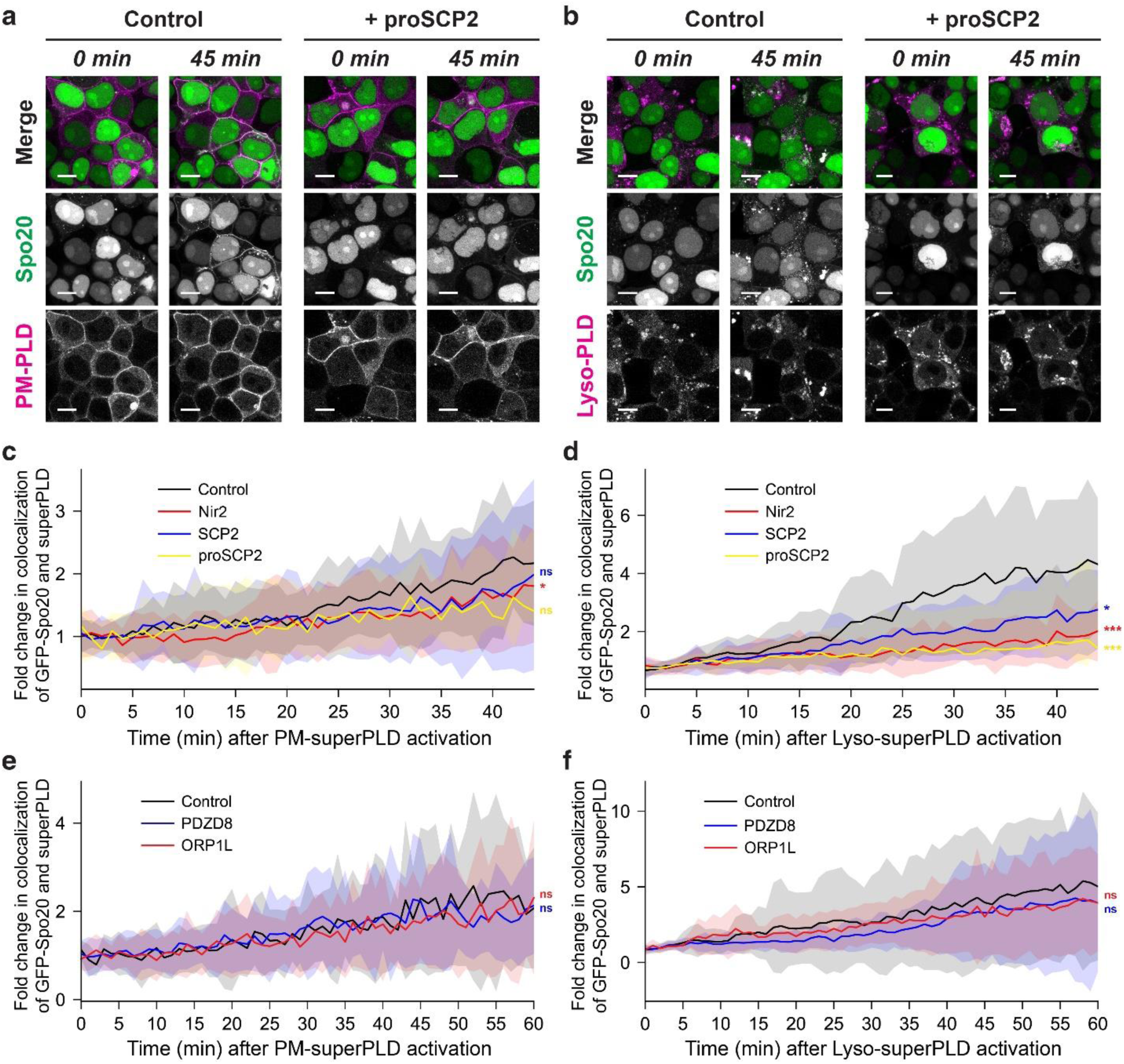
The lipid transfer protein SCP2, but not PDZD8 and ORP1L, reduces local PA enrichment. **a–b**, Confocal microscopy images of HEK 293T cells co-expressing the PA-binding probe GFP-Spo20 and superPLD targeted to the plasma membrane (a, PM) or lysosomes (b, Lyso), without (Control; left panels) or with (+ proSCP2; right panels) stable expression of proSCP2. Shown are images acquired 45 min following superPLD activation by intermittent blue light illumination (5 s per 1 min). Scale bars, 10 μm. **c–d**, Quantification of data from (a) and (b), measuring colocalization between GFP-Spo20 and superPLD targeted to plasma membrane (c, PM) or lysosomes (d, Lyso) over the course of the 45 min of superPLD activation. Each graph shows the colocalization timecourse in control HEK 293T cells (Control; black) or cells stably expressing V5-Nir2 (Nir2; red), SCP2 (SCP2; blue), or proSCP2 (proSCP2; yellow). Note that the V5 tag was omitted from SCP2 and proSCP2 to avoid potential interference with their post-translational cleavage. Statistical analysis was performed using repeated measures ANOVA (c, *n* = 20 cells, *p* = 0.0334 for Nir2, 0.3646 for SCP2, 0.0691 for proSCP2; f, *n* = 20 cells, *p* = 0.0008 for Nir2, and 0.0315 for SCP2, and 0.0002 for proSCP2). Three independent experiments were performed with similar results. **e–f**, Quantification of similar studies as (c–d) but with different lipid-transfer proteins, PDZD8 and ORP1L, measuring colocalization between GFP-Spo20 and superPLD targeted to plasma membrane (e, PM) or lysosomes (f, Lyso) over the course of 60 min following superPLD activation by intermittent blue light illumination (5 s per 1 min). Each graph shows the colocalization time course in the control HEK 293T cells (Control; black) or cells stably expressing V5-PDZD8 (PDZD8, blue) or V5-ORP1L (ORP1L, red). Imaging data for PDZD8 and ORP1L studies are shown in Fig. S2a–b. Statistical analysis was performed using repeated measures ANOVA (e: *n* = 20 cells, *p* = 0.8562 for PDZD8 and 0.5025 for ORP1L; f: *n* = 20 cells, *p* = 0.3918 for PDZD8 and 0.5803 for ORP1L). Two independent experiments were performed with similar results.

In contrast to SCP2, expression of the other two lipid-transfer proteins tested, PDZD8 and ORP1L, did not induce a decrease in PA enrichment from PA-fed membranes (**Fig. 5e–f** and **Fig. S2a–b**). Because both PDZD8 and ORP1L can bind to PA in vitro^31,32^, we examined whether superPLD-induced PA enrichment caused any changes in their subcellular localization using GFP fusions to these proteins. PDZD8-GFP was found mostly at the ER, similar to PDZD8-V5 immunofluorescence, and GFP-ORP1L showed some lysosomal association along with a cytosolic pool. It is noteworthy that the lysosomal pool of GFP-ORP1L exhibited limited colocalization with lysosome-targeted superPLD, suggesting that ORP1L and the p18/LAMTOR-targeted superPLD may reside in different microdomains (**Fig. S2c–d**). Although superPLD and the two GFP-tagged LTPs showed relatively high association with each other, we observed no significant increase in their colocalization upon superPLD recruitment to lysosomes. Altogether, these results suggest that Nir2 and SCP2 can reduce local enrichment of PA molecules at the site of PA production and that PA binding by ORP1L and PDZD8 may have other functions, as examined below.

### Lipidomics analysis reveals cells maintain PA homeostasis to counteract PA overproduction

In our Feeding-Fishing proteomics studies, several enzymes that degrade or produce PA were, respectively, either enriched on or depleted from PA-fed membranes. Notably, these enzymes displayed differential abundances between PA-fed plasma membranes and lysosomes (**Fig. S3a**). For example, all three isoforms of the lipin PA phosphatases (LPIN1/2/3) were enriched on PA-fed plasma membranes, whereas only LPIN1 was detected on PA-fed lysosomes, and at lower abundance levels. To understand the potential impact of these differences on PA metabolism and overall phospholipid homeostasis, we investigated the global changes to the phospholipidome by LC–MS-based lipidomics after superPLD-mediated membrane editing to feed PA to either the plasma membrane or lysosomes (**Fig. 6a**).

**Figure 6.**
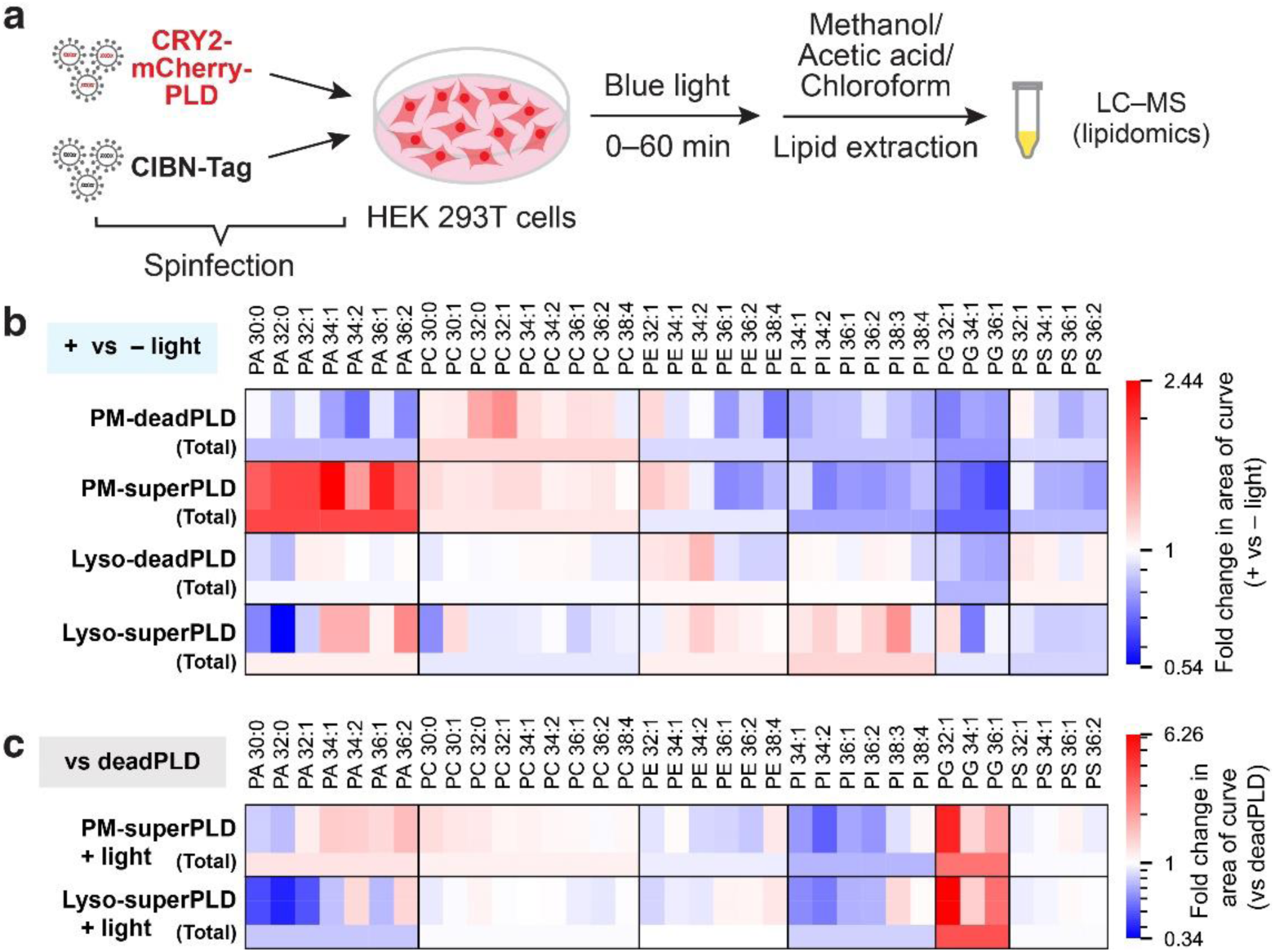
Cells maintain PA homeostasis in response to superPLD-mediated PA feeding. **a,** Schematic depiction of PA feeding coupled to lipidomics to examine changes to cellular phospholipid levels. Lentiviral spinfection was used to express superPLD or catalytically dead PLD targeted to the plasma membrane or lysosomes. 48 h post spinfection, cells were illuminated with or without intermittent blue light (5 s per 1 min) for 60 min to induce PLD recruitment to the plasma membrane or lysosomes. Cellular lipids were then extracted and analyzed by LC–MS. **b,** Heat maps depicting fold changes in individual (top) and total (bottom) levels of phospholipid species in cells after (+ light) vs. before (– light) 60-min illumination with blue light to recruit deadPLD or superPLD to either the plasma membrane (PM) or lysosomes (Lyso). **c,** Heat maps depicting fold changes in individual (top) and total (bottom) levels of phospholipid species, comparing cells expressing superPLD vs. deadPLD, after (+ light) their targeting to either the plasma membrane (PM) or lysosomes (Lyso). n=3 replicates per condition.

We found that HEK 293T cells expressing plasma membrane or lysosome-targeted superPLD exhibited light-dependent increases in PA levels after blue light-induced superPLD recruitment to the target organelle membrane (**Fig. 6b**). Such increases in PA required superPLD activity, as they were not observed upon recruitment of deadPLD to these membranes (**Fig. 6b**). Interestingly, a more pronounced PA increase was observed at the plasma membrane compared to lysosomes upon PA feeding, indicating a slower net turnover of PA molecules at the plasma membrane. Yet, the higher abundance of lipins at PA-fed plasma membranes (**Fig. S3a**) suggests an increased conversion of plasma membrane PA pools to DAG. However, the conversion of PA to DAG is bidirectional in cells (where the reverse reaction is catalyzed by DAG kinases), and the rapid turnover of DAG molecules has been observed with an estimated half-life of one to several minutes^33–35^. By contrast, the conversion of PA to CDP-DAG (and subsequently to PG and PI) is unidirectional. It is possible that the sustained buildup of PA at the plasma membrane results from increased PA flux within the bidirectional PA–DAG pathway, eventually leading to slower net PA turnover, and the increased abundance of lipins and DAG kinase eta (DGKH) on PA-fed plasma membrane could be involved in this event (**Fig. S3**). Concomitantly, the increased levels of PI in PA-fed lysosomes may be accounted for by PA flux into the PA–CDP-DAG pathway.

While PA levels increased rather modestly in superPLD-expressing cells, a striking increase was observed in PG levels (**Fig 6c**), supporting the notion that cells maintain PA homeostasis by removing excess PA molecules via the CDP-DAG pathway. This PG accumulation in superPLD-expressing cells had little light dependency (**Fig. 6b–c**), suggesting that it results from the lower but chronic PA production that occurs from background superPLD activity in the dark^18,36^. Moreover, despite superPLD consumes PC (with its preference on species with 34:1 and 36:1 acyl chains^8^) for PA production, the levels of these lipid species did not change, indicating that a substantial portion of PA produced on this timescale is routed to the Kennedy pathway as well to replenish these phospholipids (**Fig S3b**). Collectively, these lipidomics studies, along with results from imaging analysis, demonstrate that superPLD activation can indeed lead to temporally controlled local increases in cellular PA levels but that global PA levels are subject to homeostatic regulation to within a certain range to avoid excess PA accumulation.

### Lipid transfer proteins facilitate PA clearance to maintain PA homeostasis

Beyond these changes in recruitment of PA-metabolizing enzymes, the Feeding–Fishing experiments also revealed recruitment of several LTPs that could potentially be involved in restoring PA homeostasis (**Fig. 3**). Indeed, our confocal microscopy analysis in cells co-expressing superPLD and the PA-binding GFP-Spo20 probe demonstrated that forced expression of two such LTPs, Nir2 and SCP2, can diminish local enrichment of PA at the plasma membrane and lysosomal membranes (**Fig. 4–5**). To directly investigate the roles of these lipid transporters in the global regulation of PA metabolism, we performed lipidomics analysis on cells subjected to PA feeding of either the plasma membrane or lysosomes that also overexpressed one of these LTPs (**Fig. 7a**). We found that forced expression of either Nir2 or SCP2 caused a significant attenuation in the global increase of PA levels that occurs upon PA feeding of either organelle membrane (**Fig. 7b– c**). These two LTPs had a stronger effect in decreasing excess PA at the plasma membrane compared to at lysosomes, a finding that aligns with our lipidomics analysis indicating slower baseline levels of PA turnover at the plasma membrane (**Fig. 6b**). Combined with the imaging analysis, these lipidomics results suggest that Nir2 and SCP2 can facilitate PA clearance at both local and global levels.

**Figure 7.**
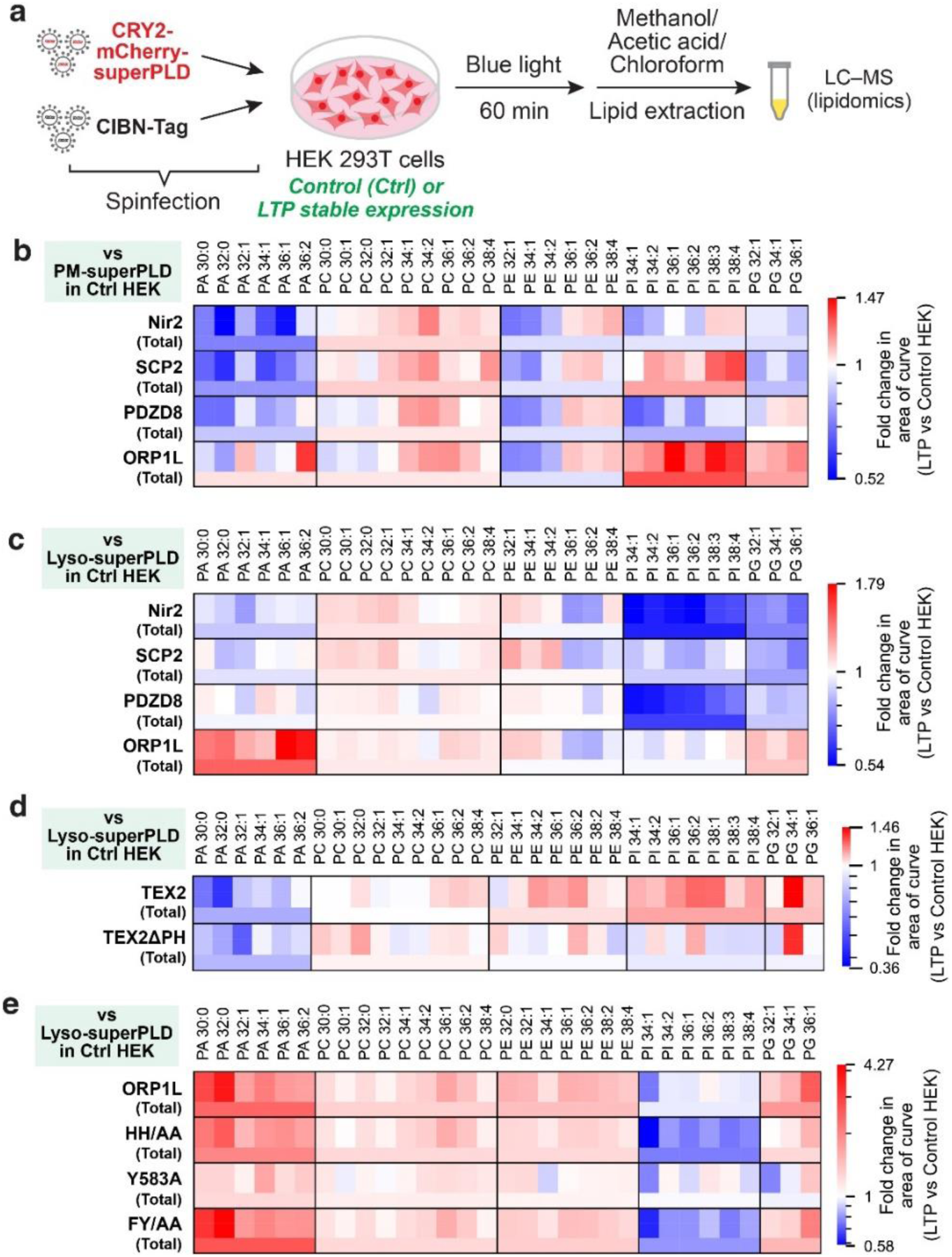
Lipid transfer proteins have divergent effects on PA metabolism. **a,** Schematic depiction of PA feeding coupled to lipidomics to examine the effects of LTPs on phospholipid homeostasis. Lentiviral spinfection was used to express superPLD targeted to either the plasma membrane or lysosomes in HEK 293T cells stably expressing an LTP. 48 h post spinfection, cells were illuminated with intermittent blue light (5 s per 1 min) for 60 min to induce superPLD recruitment. Cellular lipids were then extracted and analyzed by LC–MS. **b–c,** Heat maps showing fold changes in individual and total levels of phospholipid species in cells stably expressing V5-Nir2, V5-SCP2, PDZD8-V5, or V5-ORP1L, 60 min following recruitment of superPLD to either the plasma membrane (b) or lysosomes (c). Fold changes were calculated based on phospholipid levels in control HEK 293T cells (Ctrl) expressing empty V5 vector and otherwise treated identically. **d,** Heat maps depicting fold changes in individual and total levels of phospholipid species in cells stably expressing V5-TEX2 or V5-TEX2ΔPH (ΔPH; PH domain deleted)^40^, 60 min following recruitment of superPLD to lysosomes. Fold changes were calculated based on phospholipid levels in control HEK 293T cells (Ctrl) that were treated similarly. **e,** Heat map analysis similar to that shown in (a), but depicting fold changes in individual and total levels of phospholipid species in cells stably expressing V5-ORP1L, V5-ORP1L^H651A/H652A^ (HH/AA, a ORD domain mutant lacking PI4P-binding motif), V5-ORP1L^Y583A^ (Y583A, a ORD domain mutant lacking cholesterol-binding motif), or V5-ORP1L^F476A/Y477A^ (FY/AA, an FFAT mutant deficient in binding to VAP)^26^. n=3 replicates per condition.

We next examined the PA-binding LTPs whose overexpression did not affect GFP-Spo20 localization in our imaging assays, using the lipidomics approach. First, we found that PDZD8 overexpression led to a modest but significant decrease in PA enrichment on PA-fed plasma membranes and lysosomes (**Fig. 7b–c**). PDZD8 notably contains an SMP domain, which in other SMP domain-containing proteins can mediate glycerolipid transfer at membrane contact sites^38^. TEX2, another SMP domain-containing protein, was also identified in our Feeding–Fishing proteomics to be highly enriched on PA-fed lysosomes (**Fig. S4a**), and interestingly a similar decrease in PA enrichment was observed on PA-fed lysosomes from cells overexpressing TEX2 (**Fig. 7d**). Though PDZD8 and TEX2 share similar SMP domains and ER-anchoring hydrophobic motifs, they additionally contain different domains that mediate binding to anionic lipids and/or proteins on other membranes, i.e., C1 and PDZ domains in PDZD8^39^ and a PH domain in TEX2^40^. Overexpression of a truncated TEX2 construct lacking its PH domain (TEX2ΔPH)^40^ still induced a decrease in PA levels upon PA feeding (**Fig. 7d**), indicating that the PH domain, which can recognize several phosphoinositides in vitro^40^, is not necessary for the PA clearance. By confocal microscopy, pools of PDZD8-GFP and TEX2-GFP clustered proximal to, but not completely overlapping with, lysosome-targeted superPLD (**Fig. S4b–d**). Therefore, it is plausible that PDZD8 and TEX2 contribute indirectly to PA clearance either by mediating lipid transfer at or facilitating formation of ER–lysosome contact sites, rather than by directly binding to PA or PA-fed membranes.

Finally, in cells overexpressing ORP1L, we observed an unexpected elevation in PA levels upon PA feeding, particularly at lysosomes (**Fig. 7b–c**). This PA enrichment persisted when full-length ORP1L was replaced with ORP1L variants bearing point mutations in either the PI4P-binding site (H651A/H652A; HH/AA^26^), in a well-conserved lipid-binding motif, EQVSHHPP, located within the ORD domain^41^, or its FFAT motif (F476A/Y477A; FY/AA^26^), which mediates ER targeting via interactions with VAPA/B. However, the PA enrichment was indeed antagonized by expression of an ORP1L mutant in its ORD domain that is deficient in cholesterol binding (Y583A^42^) (**Fig. 7e**).

## Discussion

In this study, we introduce Feeding–Fishing as a general strategy to identify regulators of lipid homeostasis at the organelle level. In this approach, membrane editing is used to “feed” lipids of interest to designated organelle membranes and proximity labeling is used to “fish out” proteins recruited to those same membranes. We implemented this concept to reveal proteins and mechanisms associated with PA metabolism and transport on the cytosolic leaflets of two distinct organelle membranes: the plasma membrane and the lysosomal membrane. We subsequently analyzed the consequences of the observed enrichment of several protein hits identified in the Feeding–Fishing proteomics using a combination of confocal microscopy to visualize the subcellular localization of PA using a PA-binding probe, and LC–MS-based lipidomics to assess changes to the phospholipidome.

Our studies revealed that, following membrane editing to locally elevate PA on the plasma membrane and lysosomes, metabolic machinery is indeed recruited to deplete the excess PA, but not all PA-metabolizing pathways are called upon equally to accomplish this task. Among the several metabolic fates of PA, the enzymes involved in PA–DAG interconversion were most dynamically regulated on PA-fed membranes, whereas minor changes were observed for enzymes that mediate metabolism to LPA. The synthases that convert of PA to CDP-DAG (CDS1/2) were not reliably detected in our proteomics, likely because these are multi-pass transmembrane proteins localized in ER membranes. It would be interesting to expand the Feeding–Fishing approach to additional organelle membranes involved in PA metabolism (e.g., ER, Golgi complex, mitochondria). A limitation of our study design is the moderate background activity of superPLD that is present even without light stimulation. Light-independent PG accumulation was observed in cells expressing superPLD, indicating that this background superPLD activity had altered lipid metabolism even in this relatively short-term period (i.e., 48 h post-lentiviral transduction). Thus, future implementations of Feeding–Fishing proteomics would benefit from using LOVPLD, a newer and ultralow background optogenetic membrane editor, for PA feeding.

In addition to insights related to the contributions of different PA-metabolizing enzymes to PA homeostasis on the plasma membrane and lysosomal membrane, the Feeding–Fishing studies revealed translocation of several LTPs to PA-fed plasma membranes and lysosomes that could return excess PA and its metabolites to the ER and/or other organelles for further metabolism. Notably, both a PA-specific LTP, Nir2, and a more broad-spectrum LTP, SCP2, could reduce local PA enrichment as well as global PA levels under the conditions when PA was supplied to plasma membrane or lysosomes. The finding that SCP2 can reduce local PA enrichment suggests that this soluble LTP can mediate intracellular transport of PA, in addition to cholesterol and several other lipids that it has previously been found to transport^27,43–45^.

Unlike SCP2, Nir2 also led to reduced PI levels accompanied with PA decrease, especially when PA was fed on lysosomes. These observations are consistent with previously described functions of Nir2 as a PI–PA exchanger at ER–plasma membrane contact sites^6,46^; in this case, PA transfer from lysosomes to the ER by Nir2 would be coupled to PI transfer in the reverse direction, from the ER to lysosomes. Interestingly, a significant decrease in PI was also observed in cells overexpressing PDZD8. Although previous studies have identified PDZD8 as capable of extraction of various lipids, including PS, PA, PC, PE, ceramides, and cholesterol, in FRET-based in vitro lipid transfer assays^31,39^, its function in PI transfer — particularly in a cellular context — was not previously identified.

In contrast to Nir2, SCP2, PDZD8, and TEX2, whose overexpression antagonized the PA accumulation elicited by membrane editing, ORP1L forced expression led to an unexpected further increase in PA levels. ORP1L binds to PA^32^, but the functional relevance of such binding, and whether it is accompanied by PA extraction and interorganelle transport, remains unknown. Our structure–function studies revealed that the PA increase induced by ORP1L required the cholesterol-binding site in ORD domain but not the EQVSHHPP signature motif found in all ORP family proteins^41,47^, which is consistent with the lack of enrichment of other members of the ORP family in our Feeding–Fishing proteomics results. In other ORP family proteins, the ORD domain mediates lipid transfer by extracting the lipid from one membrane and depositing it into another^48–50^. It can have differential affinities for two different lipids whose binding sites partially overlap, and such dual affinities are critical for vectorial transport when coupled to other metabolic steps on the origin and destination membranes. The implication of the ORD domain from ORP1L in the paradoxical increase in PA seen upon ORP1L forced expression on PA-fed membranes suggests that more complex mechanisms than simple transport to the ER and subsequent metabolism, as would be predicted by analogy with other ORP proteins, might be at play. Therefore, the roles of ORP1L in PA metabolism warrant further detailed study.

Intriguingly, several other proteins related to cholesterol biosynthesis and trafficking were identified in the Feeding–Fishing proteomics. For example, SOAT1, which catalyzes the formation of fatty acid-cholesterol esters on the ER, and LIPA, which catalyzes the reverse reaction, were both enriched on PA-fed membranes. By contrast, STARD3, an LTP that mediates ER-to-lysosome cholesterol transport^51^, was depleted from PA-fed membranes. Investigation of these proteins might shed light on new and unexpected regulatory mechanisms between PA and cholesterol metabolism and homeostasis. Finally, the enrichment of select members of the bridge-like lipid transfer protein family to PA-fed membranes (VPS13B to the plasma membrane and VPS13C to the lysosome) suggests that increased bulk phospholipid flow between these organelles may occur as a result of changes to PA metabolism induced by superPLD-mediated membrane editing.

Beyond elucidating mechanisms that implicate select PA-metabolizing enzymes and transporters in mediating PA homeostasis at the organelle and cellular levels, our study is notable for the introduction of the Feeding–Fishing strategy. This approach combining membrane editing with proximity labeling is geared to reveal how targeted modifications to the lipid composition of individual organelle membranes impacts the proteomes of such membranes, information that can point to new ways that cells can sense and correct imbalances in lipid metabolism. By exploiting organelle-specific PC hydrolysis by optogenetic PLDs, we have focused here on regulators of PA metabolism at the plasma membrane and lysosomes. Yet, our optogenetic PLDs can use exogenously supplied primary alcohols in transphosphatidylation reactions to produce other phospholipids^8,18,36,52^, and a growing collection of membrane editors targeting diverse types of lipids (e.g., phosphoinositides, DAGs, sterols) is emerging^16,52–54^. The interfacing of such tools with organelle membrane proteomics using proximity labeling in Feeding–Fishing (or, conversely, Fasting–Fishing) experiments represents a powerful and generalizable strategy for elucidating mechanisms governing lipid homeostasis.

## Materials and Methods

### Plasmids and cloning

References and/or sequences of plasmids and primers used for this study are provided (**Supplementary Table 4**). Membrane-targeted TurboID and optoPLD were cloned into pCDNA5/FRT/TO for transient transfection and pCDH-CMV-MCS-EF1α-Puro for lentiviral transduction. For plasma membrane targeting, the CAAX domain of K-Ras^16^ (GKKKKKKSKTKCVIM) was fused to the C-terminus of the constructs after a linker sequence (GGSGSLYK). For lysosomal membrane targeting, the p18 domain^17^ (MGCCYSSENEDSDQDREERKLLLDPSSPPTKALNGAEPNY) followed by a linker sequence (GGRGSGSGSGSGSGSGSGSGSG) was fused to the N-terminus of the constructs.

Plasmids encoding PITPNM1 (Nir2), SCP2, PDZD8, TEX2, or OSBPL1A (ORP1L) were purchased from the DNASU Plasmid Repository, and their open reading frames were cloned into pCDH-CMV-MCS-EF1α-Puro with an optional V5 tag to generate stable cell lines overexpressing these proteins. V5-Nir2, SCP2, proSCP2 (405–547 aa of SCP2), V5-ORP1L, PDZD8-V5, and TEX2-V5 were used for analyzing their functions in PA trafficking and metabolism in live cells (note that V5 tag was omitted from SCP2 and proSCP2 to avoid potential interference with their post-translational cleavage). Nir2, PDZD8, TEX2, and ORP1L were also cloned into pCDNA3.1 along with EGFP or miRFP fluorescent proteins for localization studies in live cells.

For visualization of PA localization in live cells, a Spo20 PA-binding domain^23^ (MDNCSGSRRRDRLHVKLKSLRNKIHKQLHPNCRFDDATKTS) fused to EGFP was cloned into pCDH-CMV-MCS-EF1α-Puro.

### Mammalian cell culture and transient transfection

Cells were grown in DMEM (Corning) supplemented with 10% FBS (Corning), 1% penicillin/streptomycin (Corning), and 1 mM sodium pyruvate (Thermo Fisher) at 37 °C in a 5% CO_2_ atmosphere. For poly-L-lysine pre-treatment, cell plates were treated with 0.1 mg/mL poly-L-lysine (Sigma Aldrich; P2636) in PBS for 1 h at 37 °C, followed by triple rinses with autoclaved deionized water. For transient transfection, HEK 293T cells were transfected using Lipofectamine 2000 (Invitrogen; 11668019) following the manufacturer’s protocol. Briefly, cells were incubated in regular DMEM media containing plasmids pre-mixed with Lipofectamine 2000 (1–1.5 µg total plasmids and 3 µL Lipofectamine 2000 for cells in 35-mm dish), and the cells were incubated for 20–24 h before experiments.

### Highly efficient multi-gene expression by virus semipurification and spinfection

HEK 293TN cells were transfected using Lipofectamine 2000 or PEI MAX (Polysciences; 24765-100) following the manufacturer’s protocol for lentivirus production. Briefly, HEK 293TN cells seeded on a 6-well plate were incubated in Transfectagro (Corning) supplemented with 10% FBS containing plasmids pre-mixed with Lipofectamine 2000 or PEI MAX (0.5 µg envelope plasmid, 1 µg packaging plasmid, 1.5 µg optoPLD plasmid, and 6 µL Lipofectamine 2000 or PEI MAX per well for a 6-well plate). 8–12 h after transfection, the transfection media was replaced with regular DMEM media, and media were collected 30 h and 54 h after transfection to obtain virus-containing media. The collected virus media were concentrated by centrifugation at 100,000 g (24,000 rpm in an SW 41 Ti swinging-bucket rotor) for 90 min at 4 °C. After ultracentrifugation, the supernatant was carefully decanted, and the virus concentrate was resuspended in fresh DMEM media.

For spinfection, HEK 293T cells seeded on a 6-well plate (pre-treated with poly-L-lysine) were incubated in virus-resuspended media supplemented with 0.4 µg/mL polybrene (Millipore Sigma). The plate was centrifuged at 931 g for 2 h at 37 °C, followed by the replacement of virus-containing media with fresh DMEM media. The 6-well plate was covered with aluminum foil to keep cells in the dark, and the cells were incubated for 40–48 h before experiments.

The ratio of lentivirus-packaging (HEK 293TN) cells and transduced (HEK 293T) cells used for this study is 1.5:1 for pCDH-CRY2-mCherry-PLD constructs and 0.16:1 for all the other constructs, optimized based on the lentivirus titer of each construct.

### Setup for optogenetics experiments

A homemade light box was built by attaching four strips of dimmable, 12 V blue-LED tape light (1000Bulbs.com; 2835–60-IP65-B1203) on the inside of a Styrofoam box. For optogenetics experiments, the light box was placed inside the CO_2_ incubator using an AC Outlet Power Bank (Omars; 24,000 mAh, 80 W) as a power supply. An outlet timer (BN-LINK) was used to switch the light on and off automatically to enable 3-s intervals of blue light in every 1 min.

### Membrane-TurboID and PA-TurboID labeling and proteomics

Cells expressing optoPLD and TurboID were illuminated for 30 min with intermittent blue light (5 s pulses every min), followed by 3 min treatment with 500 µM biotin. The cells were then washed for 5 times with PBS and lysed in RIPA lysis buffer (50 mM Tris-HCl, pH 7.4, 150 mM NaCl, 1% Triton-X, 0.5% sodium deoxycholate, 0.1% SDS, 1 mM EDTA, 1x cOmplete™ Protease Inhibitor). For proteomics, the cells from all six wells in a 6-well plate (∼2 × 10^7^ cells total) were combined to prepare each sample. After sonication and centrifugation, the lysate supernatant was incubated with Pierce™ High-Capacity Streptavidin Agarose beads (Thermo Fisher Scientific; 20359) for overnight at 4 °C on a rotator. 10 µL of beads solution was used for cells per well in 6-well plate.

After overnight incubation, the streptavidin beads were washed for twice with RIPA buffer, once with 1 M KCl, once with 0.1 M Na_2_CO_3_, once with 2 M urea in 10 mM Tris-HCl, and twice with RIPA buffer. For Western blot analysis, the washed beads were boiled for 10 min in 3x Laemmli sample buffer supplemented with 25 mM biotin to elute biotinylated proteins off the beads. For proteomics, the beads were further washed twice with HEPES buffer (50 mM HEPES, 150 mM NaCl) and once with sample elution buffer (50 mM HEPES, 150 mM NaCl, 1% SDS, 0.1% triton-X, 25 mM biotin, pH 8). The washed beads were boiled for 5 min in sample elution buffer for protein elution. The elution process was repeated for three times, each time replacing the elution buffer, to increase the protein yield. The samples were then quantified by BCA assay and submitted to Proteomics and Metabolomics Facility at Cornell University for TMT-labeling and proteomics analysis. Four samples in triplicate were grouped for the labeling and analysis.

### Analysis of proteomics data

The proteomics data contained the abundance of different proteins detected in each sample. We screened for proteins that showed significant increase (fold change > 1 and fold change *p*-value < 0.05) or decrease (fold change < 1 and fold change *p*-value < 0.05) based on protein abundance ratio. Gene ontology enrichment analysis was performed to identify proteins linked to biological process related to lipids. Corneocyte proteins, keratin and filaggrin, which were considered to be detected due to contamination, were removed from the search.

### Timecourse imaging by confocal microscopy

Cells were seeded on 35-mm glass-bottom imaging dishes (Matsunami Glass). Images were acquired every 1 min for 1 h at 37 °C using Zeiss Zen Blue 2.3 on a Zeiss LSM 800 confocal laser scanning microscope equipped with Plan Apochromat objectives (40X 1.4 NA) and two GaAsP PMT detectors. Solid-state lasers (488, 561, and 640 nm) were used to excite GFP, mCherry, and miRFP, respectively, and the 488 nm laser irradiation also served as a stimulus for activating optoPLD recruitment to the plasma membrane, ER, or lysosomes.

### Quantification of PA enrichment on the membranes

HEK 293T cells co-expressing a PA-binding probe (EGFP-Spo20), superPLD (CRY2-mCherry-superPLD^med^, derived from superPLD clone 1-12^18^) targeted to different organelle membranes, and the indicated lipid-transfer protein were imaged as described above. Colocalization analysis between PA-binding probe and superPLD was carried out on ImageJ as follows. First, an ROI was drawn around each cell expressing both constructs. Second, for each ROI, the mCherry signal of the superPLD was used to generate a binary mask. Third, the ratio of the EGFP signal from the PA-binding probe found in the mask, as compared to the total signal, was calculated to obtain the colocalization ratio. Finally, the fold-change in this colocalization ratio was tracked over the course of the time series and plotted with the use of matplotlib in Python. Statistical analysis was performed by Repeated Measures ANOVA.

### Immunofluorescence imaging

Immunofluorescence imaging was performed as described previously with some modifications^8^. Briefly, cells were fixed in 4% formaldehyde for 10 min at room temperature, rinsed three times with PBS, permeabilized with 0.5% Triton X-100 in PBS for 15 min at room temperature, and blocked with 1% BSA and 0.1% Tween-20 in PBS (blocking buffer) for 30 min. The cells were then treated with a 1:100 dilution of anti-V5 antibody (Bio-Rad; MCA1360GA) in blocking buffer for 1 h at room temperature and rinsed three times with 0.1% Tween-20 in PBS (PBS-T).

For immunofluorescence imaging after TurboID labeling, the cells were treated with a mixture of 1:100 dilution of streptavidin–Alexa Fluor 488 (Thermo; S11223) and 1:1000 dilution of anti-mouse–Alexa Fluor 647 (Invitrogen; A31571) in blocking buffer for 1 h at room temperature and rinsed three times with PBS-T. For immunostaining lipid-transfer proteins, the cells were treated with a 1:1000 dilution of anti-mouse–Alexa Fluor 488 antibody conjugate (Invitrogen; A21202) in blocking buffer for 1 h at room temperature and rinsed three times with PBS-T. After the final rinse, the cells were incubated in PBS-T for 5 min and stored in PBS supplemented with 1 µg/mL DAPI. Image acquisition by laser-scanning confocal microscopy was performed as described above by using solid-state lasers (405, 488, 561, and 640 nm) to excite DAPI, Alexa Fluor 488, mCherry, and Alexa Fluor 647, respectively.

### Lipidomics analysis by LC–MS

HEK 293T cells expressing the indicated constructs were illuminated for 30 min with intermittent blue light (5 s pulses every min), rinsed once in PBS, and the cellular lipids were extracted by Bligh and Dyer method as reported previously^8^. Briefly, 250 μL of methanol was added to cells in 35-mm dish on ice. Followed by the addition of 125 µL of 20 mM acetic acid and 100 µL of PBS, the cells were scraped off and transferred into a 1.5-mL centrifuge tube. After addition of 500 µL of chloroform, the tube was shaken vigorously for 3 min and centrifuged for 1 min at 10,000 g. The bottom organic layer was transferred into a new tube and dried under a stream of N_2_ gas. The resulting lipid film was dissolved in 150 µL of chloroform and subjected to high-resolution LC–MS analysis.

LC–MS measurement was performed on an Agilent 6230 electrospray ionization–time-of-flight MS coupled to an Agilent 1260 HPLC equipped with a Luna 3 µm Silica LC Column (Phenomenex; 50 × 2 mm) using a binary gradient elution system where solvent A was chloroform/methanol/ammonium hydroxide (85:15:0.5) and solvent B was chloroform/methanol/water/ammonium hydroxide (60:34:5:0.5). Separation was achieved using a linear gradient from 100% A to 100% B over 10 min. Phospholipid species were detected using an Agilent Jet Stream source operating in positive or negative mode, acquiring in extended dynamic range from m/z 100–1700 at one spectrum per second; gas temperature: 325 °C; drying gas 12 L/min; nebulizer: 35 psig; fragmentor 300 V (for positive mode) and 250 V (for negative mode); sheath gas flow 12 L/min; Vcap 3000 V; nozzle voltage 500 V.

The LC–MS data were analyzed on MassHunter Quantitative Analysis Software using the “Find compounds by formula” tool. The search parameters were set as follows: Source of formulas to confirm, Database/Library provided as **Supplementary Table 5**; Matches per formula, 1 (Automatically increase for isomeric compounds); Values to match, Mass and retention time (retention time required); Match tolerance, Masses +/– 20 ppm and Retention times +/– 0.200 minutes, Expansion of values for chromatogram extraction, m/z +/– 20 ppm and retention time – 0.500 + 1.00 minutes; Positive ion charge carriers, +H; Negative ion charge carriers, -H.

## Supplementary material

Supplementary Table 1. Results from Feeding–Fishing proteomics on plasma membrane.

Supplementary Table 2. Results from Feeding–Fishing proteomics on lysosomes.

Supplementary Table 3. Summary of Feeding–Fishing hits related to lipid biology.

Supplementary Table 4. List of plasmids and primers used in this study.

Supplementary Table 5. List of phospholipid mass and retention time.

## Acknowledgments

J.M.B. acknowledges support from the National Institutes of Health (R01GM151682). R.T. was supported by Honjo International, Funai Overseas, Cornell fellowships. We acknowledge Cornell University Proteomics and Metabolomics Facility for their support in designing and analyzing proteomics study; specifically, we thank Elizabeth Anderson for TMT labeling of proteomics samples, Dr. Qin Fu for data analysis, and Dr. Sheng Zhang for guidance.

## Author contributions

R.T. and J.M.B. conceived of the project, designed experiments, interpreted the results, and wrote the manuscript. R.T. carried out all experiments and data analysis.

---

**Figure S1.**
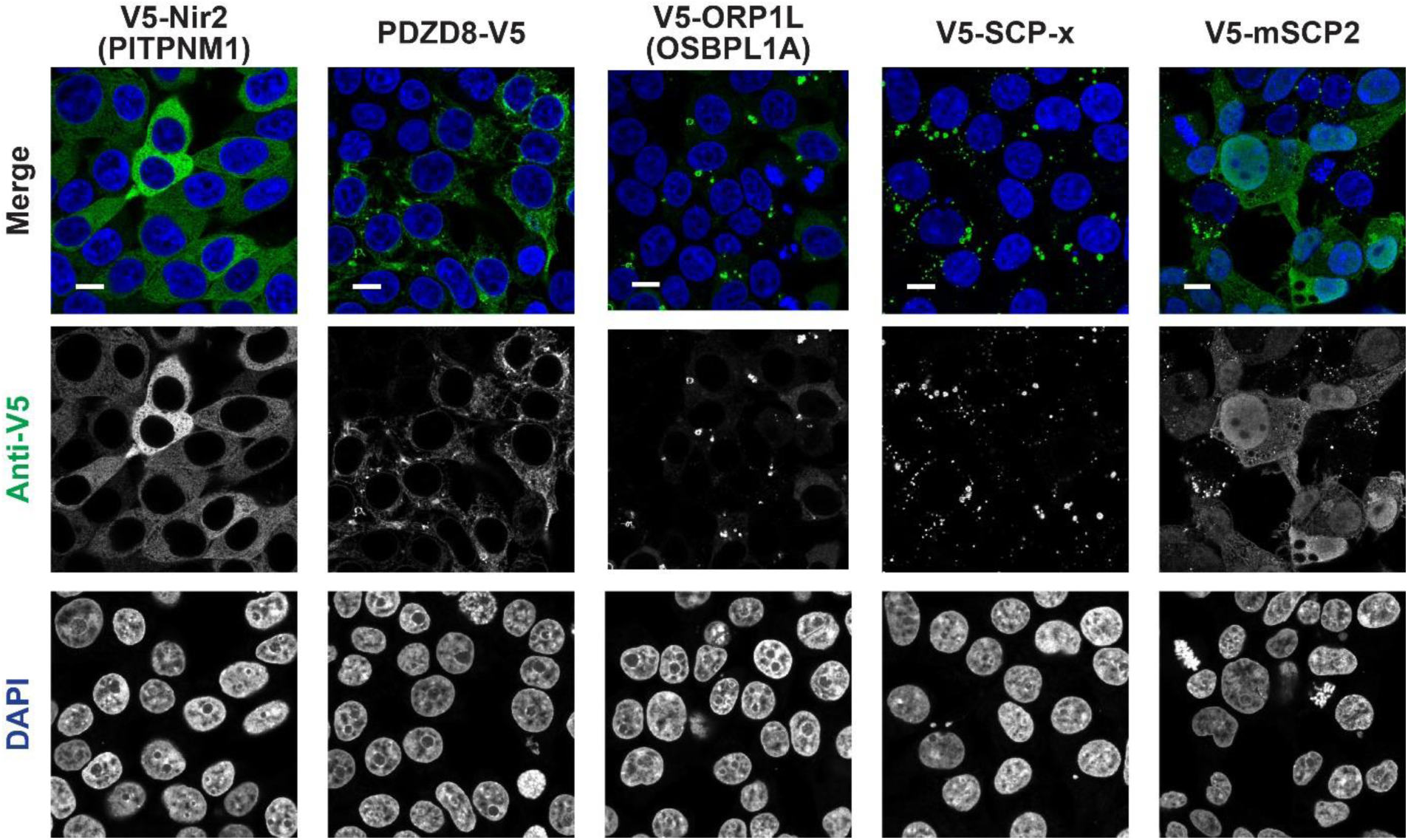
Stable expression of lipid-transfer proteins in HEK 293T cells. Confocal images of HEK 293T cells stably expressing V5-tagged Nir2 (PITPNM1), PDZD8, ORP1L (OSBPL1A), full-length SCP2 (SCP-x), or the mature form of SCP2 (mSCP2). Scale bars, 10 μm.

**Figure S2.**
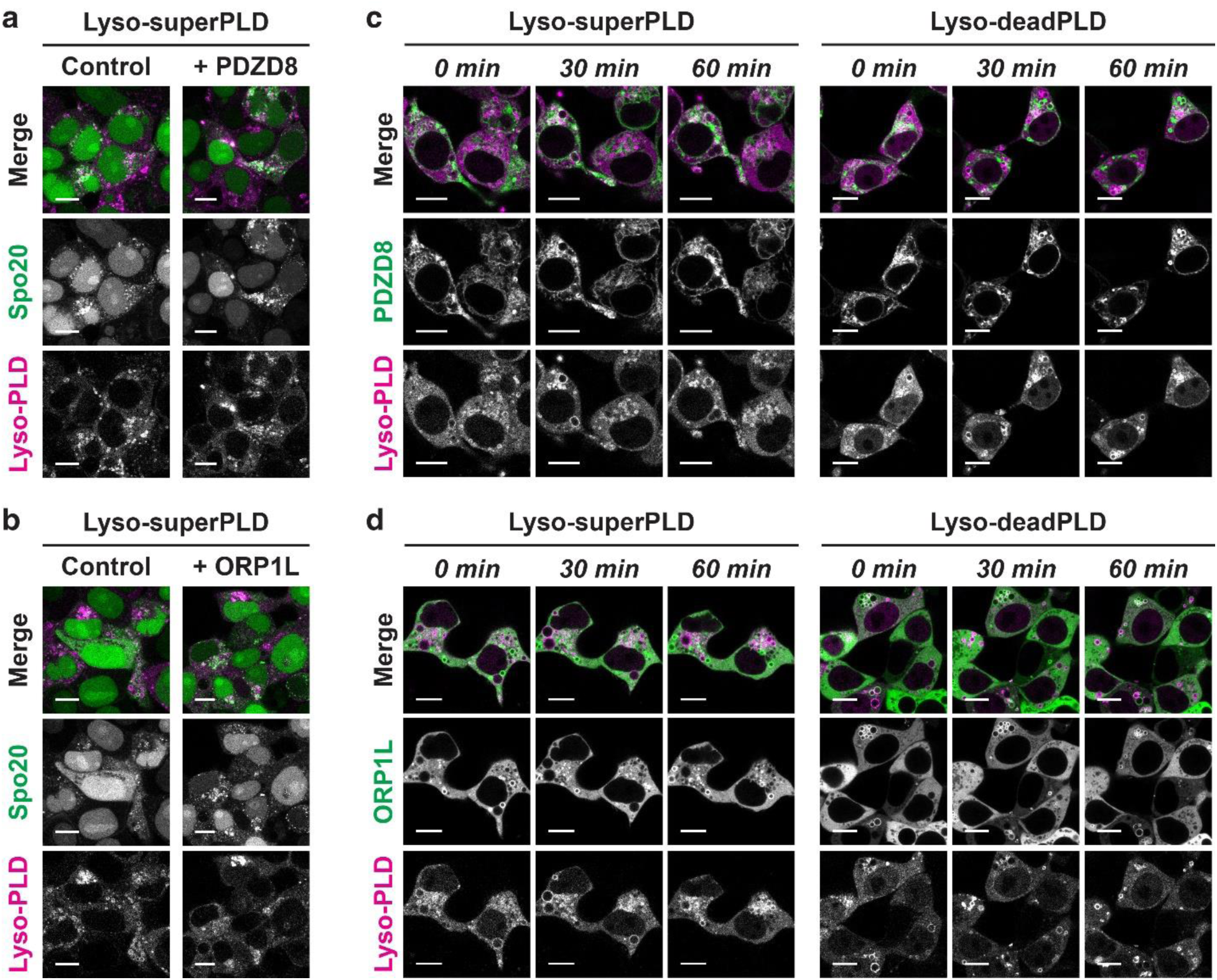
PDZD8 and ORP1L did not induce reduction of PA enrichment or recruitment to PA-enriched membranes. **a–b**, Confocal microscopy images of HEK 293T cells co-expressing the PA-binding probe GFP-Spo20 and superPLD targeted to lysosomes (Lyso), without or with stable expression of PDZD8-V5 (a) or V5-ORP1L (b). Shown are images acquired 60 min following superPLD activation by intermittent blue light illumination (5 s per 1 min). **c–d**, Confocal microscopy images of HEK 293T cells co-expressing PDZD8-EGFP (c) or EGFP-ORP1L (d) and superPLD or deadPLD targeted to lysosomes (Lyso). Images acquired 0, 30, and 60 min following recruitment of superPLD or deadPLD induced by intermittent blue light illumination (5 s per 1 min) are shown. Scale bars, 10 μm.

**Figure S3.**
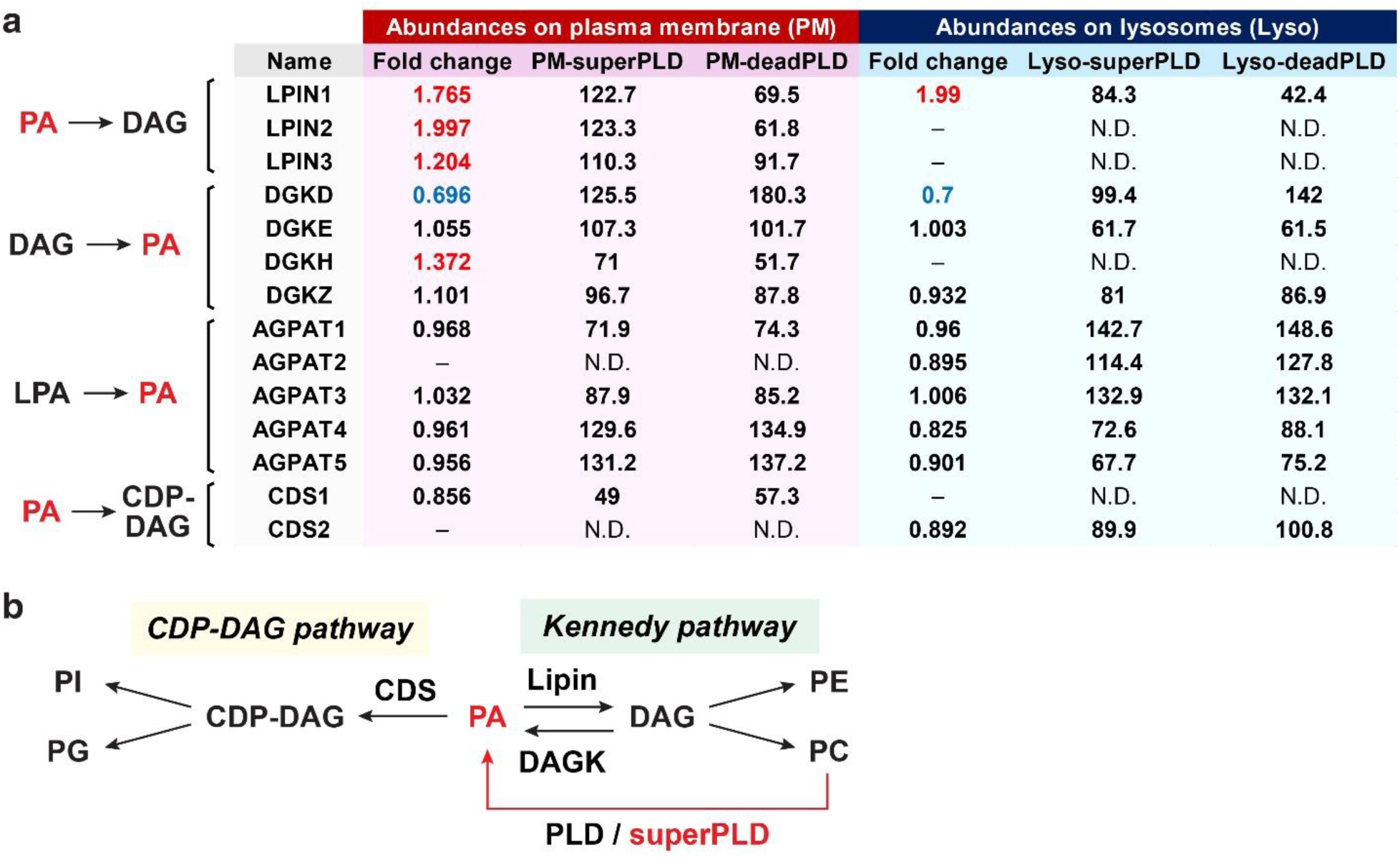
List of PA-related lipid-modifying enzymes detected in Feeding–Fishing proteomics studies. **a,** Differential enrichment and depletion of enzymes that mediate PA synthesis and degradation on PA-fed plasma membranes (left) and lysosomes (right). Shown in the table are fold changes in protein abundance between the PA-fed (superPLD-recruited) and the negative control (deadPLD-recruited) membranes and the mean abundance values for each condition (n=3). The fold changes determined to be statistically significant (abundance ratio *p*-value < 0.05) are colored in red, whereas those with significant depletion are colored in blue. Enzymes that mediate conversion of PA to LPA (e.g., PLA2s^37^) were not detected. **b,** The CDP-DAG pathway and Kennedy pathway are two major pathways by which PA is converted to other phospholipids. CDP-DAG, cytidine diphosphate diacylglycerol; DAG, diacylglycerol; LPA, lysophosphatidic acid; PA, phosphatidic acid; PC, phosphatidylcholine; PE, phosphatidylethanolamine; PG, phosphatidylglycerol; PI, phosphatidylinositol.

**Figure S4.**
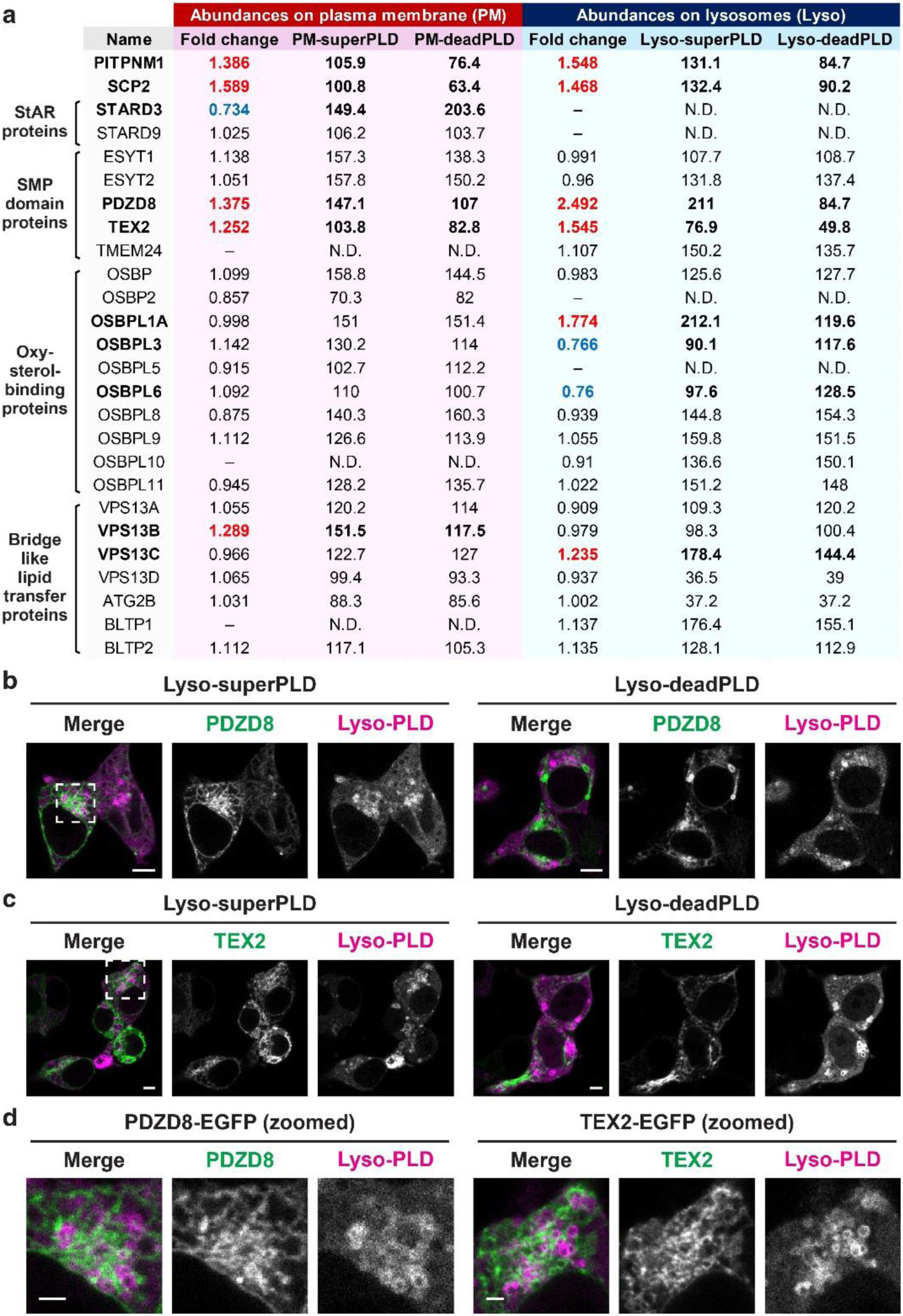
Lipid transfer proteins may directly or indirectly regulate PA metabolism. **a,** List of lipid transfer proteins exhibiting differential enrichment on or depletion from PA-fed plasma membranes (left) and lysosomes (right). Shown in the table are fold changes in protein abundance between the PA-fed (superPLD-recruited) and the negative control (deadPLD-recruited) membranes and the mean abundance values for each condition (n=3). The fold changes determined to be statistically significant (abundance ratio *p*-value < 0.05) are colored in red, whereas those with significant depletion are colored in blue. **b–c**, Confocal microscopy images of HEK 293T cells expressing PDZD8-EGFP (b) or TEX2-EGFP (c), with co-expression of superPLD or deadPLD targeted to lysosomes (Lyso). Images acquired 60 min following superPLD/deadPLD recruitment by intermittent blue light illumination (5 s per 1 min) are shown. Scale bars, 5 μm. **d**, Zoomed-in images of b–c for the areas marked with the dashed rectangles. Scale bars, 2 μm.

